# Impairments in sensory-motor gating and information processing in a mouse model of *Ehmt1* haploinsufficiency

**DOI:** 10.1101/257626

**Authors:** Brittany A. Davis, François David, Ciara O’Regan, Manal A. Adam, Adrian J. Harwood, Vincenzo Crunelli, Anthony R. Isles

## Abstract

Regulators of chromatin dynamics and transcription are increasingly implicated in the aetiology of neurodevelopmental disorders (NDDs). Haploinsufficiency of *EHMT1*, encoding a histone methyl-transferase, is associated with several NDDs, including Kleefstra syndrome, developmental delay and autism spectrum disorder. Using a mouse model of *Ehmt1* haploinsufficiency (*Ehmt1*^D6Cre/+^), we examined a number of brain and behavioural endophenotypes of relevance to NDDs. Specifically, we show that *Ehmt1*^D6Cre/+^ mice have deficits in information processing, evidenced by abnormal sensory-motor gating, a complete absence of object recognition memory and a reduced magnitude of auditory evoked potentials in both paired-pulse inhibition and mismatch negativity (MMN). The electrophysiological experiments show that differences in magnitude response to auditory stimulus were associated with marked reductions in total and evoked beta- and gamma-band oscillatory activity, as well as significant reductions in phase synchronisation. The pattern of electrophysiological deficits in *Ehmt1*^D6Cre/+^ matches those seen in control mice following administration of the selective NMDA-R antagonist, ketamine. This, coupled with reduction of *Grin1* mRNA expression in *Ehmt1*^D6Cre/+^ hippocampus, suggests that *Ehmt1* haploinsufficiency may lead to disruption in NMDA-R. Taken together, these data indicate that reduced *Ehmt1* dosage during forebrain development leads to abnormal circuitry formation, which in turn results in profound information processing deficits. Such information processing deficits are likely paramount to our understanding of the cognitive and neurological dysfunctions shared across the NDDs associated with *EHMT1* haploinsufficiency.

## INTRODUCTION

Post-translational modifiers of histone proteins influence chromatin dynamics and transcriptional regulation throughout development and are essential for the highly choreographed processes of lineage commitment and differentiation during neurodevelopment (Tyssowski et al., 2014, Hirabayashi and Gotoh, 2010). Perhaps unsurprisingly, exome sequencing studies and pathway analyses of genome wide association studies implicate genes encoding chromatin and transcriptional regulators in the aetiology of autism spectrum disorders (RK et al., 2017, De Rubeis et al., 2014), schizophrenia (Network and Pathway Analysis Subgroup of Psychiatric Genomics, 2015) and severe developmental disorders (Singh et al., 2016, Deciphering Developmental Disorders, 2017). One such gene, implicated in several neurodevelopmental and neuropsychiatric disorders (Talkowski et al., 2012), is *EHMT1*, which encodes the histone H3 lysine 9 mono- and di-methyltransferase G9a-like protein (GLP).

Haploinsufficiency of *EHMT1* is the primary cause of the 9q34 subtelomeric-deletion syndrome, also known as Kleefstra Syndrome (Kleefstra et al., 2006, Kleefstra et al., 2005), a condition associated with intellectual disabilities, epilepsy, childhood hypotonia, facial dysmorphism, microcephaly, delays in reaching developmental milestones and behavioural problems such as aggressive outbursts and hypoactivity. Furthermore, analysis of Copy Number Variants (CNVs) (Cooper et al., 2011) and a large exome sequencing study (Deciphering Developmental Disorders, 2017) have linked *de novo* mutations affecting *EHMT1* to severe developmental delay more generally. Finally, CNVs spanning *EHMT1* have also been associated with autism spectrum disorder (O’Roak et al., 2012) and schizophrenia (Kirov et al., 2012).

The importance of *Ehmt1* in brain function is supported by data from animal models that demonstrate a range of behavior changes reminiscent of neurodevelopmental disorders including exploration and/or anxiety phenotypes (Balemans et al., 2010, Schaefer et al., 2009), and abnormal learning and memory (Iacono et al., 2018, Benevento et al., 2017, Balemans et al., 2013, Kramer et al., 2011). More recently, studies using rodent neuronal cultures and *ex vivo* slices (Benevento et al., 2016), and human iPSCs (Frega et al., 2019), have shown that appropriate expression of *EHMT1* is required for the correct establishment and function neuronal networks. In human iPSCs, this neuronal network dysfunction is driven by the abnormal expression of *GRIN1* expression and enhanced NMDA-R signaling (Frega et al., 2019).

Here we explore endophenotypes of relevance to psychiatric problems seen in those carrying mutation of *EHMT1*, using a mouse model of *Ehmt1* haploinsufficiency (*Ehmt1*^D6Cre/+^ mice). In order to reduce anatomical complexity and allow a more precise focus on the impact of *Ehmt1* haploinsufficiency during development, on later behavioural and neurophysiological parameters, we generated a forebrain-specific *Ehmt1* knockout mouse. Cre recombination was driven under the D6 promoter of the *Dach1* gene limiting *Ehmt1* heterozygous deletion to the forebrain, starting at embryonic stage 10.5 in the cortical vesicles (van den Bout et al., 2002, Machon et al., 2002). Specifically, we find deficits in sensory motor-gating and novel object recognition, and decreased anxiety. We then build upon recent *in vitro* and *ex vivo* evidence of abnormal neuronal networks and show a reduction in the magnitude of auditory evoked potentials in paired-pulse inhibition and mismatch negativity, providing *in vivo* electrophysiological evidence of an impairment in information processing in the *Ehmt1*^D6Cre/+^ mice. Gene expression data and pharmacological manipulation support the general idea that abnormal NMDA-R signalling in *Ehmt1*^D6Cre/+^ adult mouse contributing to the sensory-motor gating and information processing deficits.

## MATERIALS AND METHODS

### Animals

All procedures were conducted in accordance with the requirements of the UK Animals (Scientific Procedures) Act 1986, with additional ethical approval at Cardiff University.

In order to generate experimental cohorts, *Ehmt1*^fl/fl^ (Schaefer et al., 2009) male studs were paired in trios with one homozygous females carrying two copies of the Tg(Dach1-cre)1Krs/Kctt Cre transgene, maintained on a F1(C57BL/6J x CBA/Ca) background; and one wild-type F1(C57BL/6J x CBA/Ca) female. The *Ehmt1*^D6Cre/+^ mouse model was used in order to limit the effects of the deletion to the forebrain and hippocampus only and confounding effects of the non-CNS phenotypes, such as obesity (Balemans et al., 2010). Experimental cohorts were reared together and then weaned into mixed cages (2-5per cage) of *Ehmt1*^D6cre*/+*^ (experimental line) and *Ehmt1*^fl/+^ mice (control line). All experimental subjects were male mice, and aged between 4-6 months during behavioural testing. A subset of the behavioural cohort were subsequently used in the electrophysiology experiments (7-8 months). Animals were housed 12 hour light/12 hour dark (lights on at 7am), and standard laboratory diet and water were available *ad libitum* throughout testing. Experimenters were blind to the genotype of animals during behavioural testing.

### Behaviour

All animals were initially subject to sensory-motor gating testing (*Ehmt1*^fl/+^, n=25; *Ehmt1*^D6cre*/+*^ mice, n=31). A subset of these were then subsequently tested on the rotor-rod (*Ehmt1*^fl/+^, n=9; *Ehmt1*^D6cre*/+*^ mice, n=18) and then elevated plus maze and open field. The remainder of the animals were subsequently tested on the novel object recognition memory test (*Ehmt1*^fl/+^ mice, n=16; *Ehmt1*^D6cre*/+*^ mice, n=13).

#### Sensory-motor gating

Acoustic startle response (ASR) and prepulse inhibition (PPI) of the startle response were monitored using a SR-Lab apparatus (San Diego Instruments, U.S.A) modified for use in mice, according to published methods (McNamara et al., 2016). Briefly, animals were placed in a Perspex tube (internal diameter 35mm) and white noise stimuli were presented via a speaker. The amount of movement relayed as their startle response was measured as a piezoelectric measure converted to arbitrary startle units. The measurement used was the maximum startle (Vmax). Due to the effect of weight on this reflex movement measurement, all data was normalised for individual body weight. Pulse-alone trials consisted of a 40ms 120dB startle stimulus and a prepulse trial consisted of a 20ms prepulse at 4, 8, or 16db above background and a 40ms 120db startle stimulus, 70ms after the prepulse. The stimuli were presented in a pseudorandom manner every 15s. Whole body startle responses were recorded as the average startle during a 65ms window timed from the onset of the startle pulse.

Percentage PPI score for each trial was calculated:

[%PPI=100 × (ASRstartle pulse alone - ASRprepulse + startle pulse)/ASRstartle pulse alone].

#### Rotarod testing

A rotarod task (Ugo Basile, Italy) was used to assess motor learning and co-ordination. This consisted of a rotating rod 30 mm in diameter, with five separated chambers 57 mm in width, with a rod elevation of 160 mm. Motor learning was assessed across six rotarod sessions, one morning session and one evening session, on three consecutive days. The rod speed accelerated incrementally from 5-50rpm across the 300sec session. The main measure during training was latency to first fall. However, if the mice fell, they were continuously replaced on the rotating rod, until the full 300sec-session was over in order to prevent any confounds from arising from overall differences in time spent on the rotarod across sessions. In a separate test session on day four, the mice were given one 300-sec session at 10rpm, 20rpm, 30rpm, 40rpm and 50rpm consecutively in one morning session in order to assess motor coordination. Again the latency to first fall was recorded for each animal at each speed.

#### Novel object recognition memory

The NOR test arena was a square 30cm x 30cm with 30cm high, white Perspex walls. Four different, non-displaceable objects were used. All objects were white and selected for their equal appeal and available in triplicate to avoid the use of olfactory cues. In the habituation phase, 24hr prior to the task, each subject was allowed to explore the arena for 10min in the absence of objects. In the acquisition phase, the subject was returned to the arena containing 2 identical sample objects (A, A’) and given 10min to explore. After a retention phase, consisting of 15min or 90min, the subject was returned to the arena with two objects, one identical to the sample and the other novel (A, B). During both the familiarization and test phase, objects were located adjacent corners of the arena. The location of the novel object was counterbalanced. To prevent coercion to explore the objects, the subject was released in a third corner. Objects were cleaned with 70% ethanol wipes between sessions and object selection was randomized.

The main measure used was the Recognition Index (RI), indicating whether the animal investigated the novel object more than chance. This was calculated by the percentage of time spent investigating the novel object relative to the total object investigation [RI= T_N_/(T_N_+T_F_) x 100]. An RI significantly above chance or 50% indicates recognition of novelty and an RI equal to or below 50% indicates either no preference for the novelty or no recognition of novelty. Other parameters recorded were:overall time spent with each object, and frequency of visits to the zones containing an object. Data was collected in 1-min time bins across the 10-min session by a camera linked to a computer with Ethovision Observer software (Noldus, Nottingham, UK).

### Electrophysiology

Adult male *Ehmt1*^D6Cre/+^ mice (*n*=7) and *Ehmt1*^fl/+^ control cage-mates (*n*=7) at 6-7 months of age were anesthetized with 2% isoflurane for stereotaxic electrode implantation. The electrode configuration used two bilateral frontal electrodes, one monopolar and one bipolar (2.7-mm anterior, 1.5-mm lateral, 1.2-deep relative to bregma); two bilateral hippocampal electrodes, one monopolar and one bipolar (2.7 mm posterior, 3-mm lateral, 2.2-mm deep relative to bregma); and one bipolar electrode in the auditory cortex (2.7-mm posterior, 4-mm lateral, 1.1-deep relative to bregma) as has been previously reported (Ehrlichman et al., 2008, Siegel et al., 2003) (for further details see Supplementary Figure 1). Due to animal loss, a subset of these was used in the MMN study; *Ehmt1*^D6Cre/+^ mice (*n*=5) and *Ehmt1*^fl/+^ control cage-mates (*n*=5).

ERPs were obtained by averaging EEG traces centred at time 0 and 500msec to 0μV, respectively. For each trial, power was calculated by using a complex Morlet’s wavelets w(t, f0) (Kronland-Martinet et al., 1987). The script used can be found at https://www.physics.lancs.ac.uk/research/nbmphysics/diats/tfr/. The wavelets have a Gaussian shape in the time domain (SD σt) and in the frequency domain (SD σt) around its central frequency *f*0: w(t, f0)=A * exp(-t2/2 σt2) * exp(2iπf0t) with σf=1/πσt. The wavelet family we used was defined by f0/σf=1, with f0 ranging from 0.5 to 100Hz in logarithmically distributed frequency steps (for full details see Supplementary Information - Materials and Methods).

Auditory stimuli were generated using Spike2, version 7.2 and a Power1401 interface (CED, UK). Auditory stimulus was delivered with speakers positioned directly in front of each recording cage. Each mouse received an auditory, paired-pulse session and two sessions of mismatch negativity (MMN).

#### Paired pulse

Following Halene *et al*. (Halene et al., 2009), each mouse received an auditory, paired-pulse session in which a single tone was presented at 1500Hz and 90dB (S1) followed by a 500ms intra-trial interval and a second tone at 1500Hz and 90dB (S2). The tones were sinusoidal and 10ms in duration. The inter-pulse interval between the two tones was 10-s. Each mouse received 1250 paired-pulse trials per recording session.

#### Mismatch negativity

The mice also received two sessions of the MMN protocol. Similar to Ehrlichman *et al*. (Ehrlichman et al., 2008), the mice received 24 ‘standard’ tones at 90dB and 1500Hz and one ‘deviant’ tone at 90dB and 2000Hz. All tones were sinusoidal and 10ms in duration and the intra-pulse interval between the 25 tones was 500-ms, while the inter-trial interval was 5-s. Each mouse was recorded for 360 trials in each of two sessions. In one session 10mg/kg of Ketamine was administered and in the second session an equal volume of saline was administered. The dosage of ketamine was chosen based on previous work (Ehrlichman et al., 2008, Siegel et al., 2003). The within group design was counterbalanced by genotype and the order in which ketamine or saline sessions were administered. All recordings took place 5-mins after intraperitoneal injections of either 10mg/kg ketamine or the volume equivalent dose of saline. The waveform channels were filtered between 1 and 500Hz.

### Gene Expression

RNA was extracted from macrodissected hippocampi using the RNeasy micro kit (Qiagen) following the manufacturers instructions. A 96 well RT^2^ Custom Profiler PCR Array (CAPM12608, Qiagen) for mice was used. The custom genes list was generated based on GLP targets identified in the literature and tissue-specific relevance. For the qPCR array 1μg of total RNA was used and manufacture’s instructions were followed, for cDNA synthesis—RT^2^ First Strand Kit (Qiagen) and for the RT-PCR reaction—RT^2^ SYBR Green ROX qPCR Matermix (Qiagen). The average Ct values across the three housekeeping genes *B2m, B-actin* and *Gapdh* were used as endogenous controls. ΔCt values were generated by normalizing to the geometric mean of the Ct values for these three housekeeping genes. All individual reactions were carried out in triplicate. Real time qPCR data was visualized using the ΔΔCt method (Livak and Schmittgen, 2001).

### Statistics

All data were analyzed using SPSS 20 (SPSS, USA). The statistical differences between groups were analysed using independent samples t-tests, ANOVAs, or where appropriate Repeated Measures ANOVA (RP-ANOVAs). The main between-subject factor was GENOTYPE (*Ehmt1*^*fl/+*^ controls or *Ehmt1*^*D6Cre/+*^). The following within-subject factors were also analysed: TRIAL (Startle trial); PREPULSE (4, 8 or 16dB); SESSION and SPEED (rotarod); PULSE (standard or deviant in the MMN). To check for normal distribution, Mauchly’s test of sphericity of the covariance matrix or Levenes test for equality of variances were applied. The more conservative Huynh-Feldt corrections, with adjusted degrees of freedom are reported in cases in which test assumptions were violated. Non-parametric analyses were as follows. For the NOR task, the binomial distribution one sample Kolmogorov Smirnov (KS) test was applied to determine whether average RIs or SIs were significantly above chance (above 50%). For analysis of N1 amplitude, Wilcoxon rank-sum test was used. Statistical tests on the electrophysiological time-frequency data were performed using the permutation method with <1,000 iterations (Westfall and Young, 1993) (see Supplementary Information – Materials and Methods for full details). For all comparisons, alpha value was set at *p*<0.05.

## RESULTS

### Impaired sensory-motor gating in *Ehmt1*^D6Cre/+^ mouse

The acoustic startle response and pre-pulse inhibition (PPI) of this response, were used to examine sensory-motor function in the *Ehmt1*^D6Cre/+^ mice. Over the course of six consecutive auditory pulse (120dB) startle trials, *Ehmt1*^D6Cre/+^ mice had, on average, twice the startle response compared to *Ehmt1*^fl/+^ mice (Figure 1A; ANOVA, main effect of GENOTYPE, F_1,53_=6.86, *p*=0.01). A significant interaction between GENOTYPE and TRIAL (F_3.87,201.0_=2.48, *p*=0.03) indicated different patterns of startle reactivity and habituation relative to *Ehmt1*^fl/+^ mice. *Post hoc* analysis revealed that there was an equivalent startle response to the initial trial between genotypes (*p=*0.68), but on average *Ehmt1*^D6Cre/+^ mice showed a significantly enhanced startle response in all consecutive trials (trial 2, *p=*0.05; 3, *p=*0.01; 4, *p=*0.001; 5, *p=*0.01; and 6, *p=*0.02).

**Figure 1.**
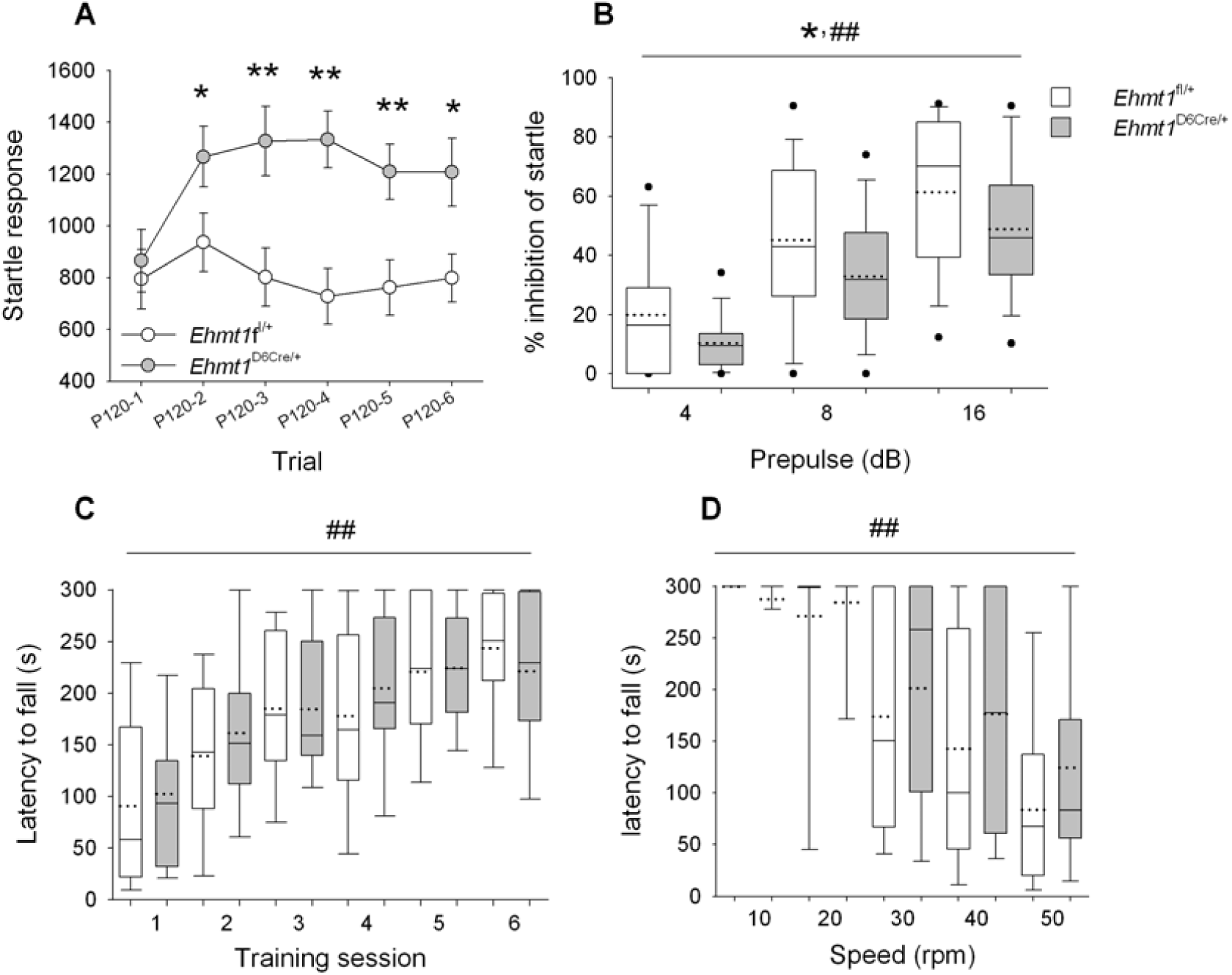
Sensory-motor gating & motor learning and coordination. **A**) The startle response after 6 consecutive 120dB pulse stimulations show *Ehmt1*^D6Cre/+^ mice (n=31) had a significantly higher overall maximum startle response in all trials. **B**) Prepulse inhibition of the startle response generally increased with the increasing volume of the prepulse as expected. However, *Ehmt1*^D6Cre/+^ displayed a 20-40% reduction in PPI of startle relative to *Ehmt1*^Fl/+^ mice (n=25) **C)** Both *Ehmt1*^Fl/+^ (n=9) and *Ehmt1*^D6Cre/+^ mice (n=18) improved motor ability on the rotarod as evidenced by an increase in latency to fall across 6 training sessions. There was no also difference in genotype during training **D)** In a probe test where rotarod speed accelerated throughout the session, there was a general decrease in the latency to fall as the speed increased, but again no difference between genotypes. Data are mean ±SEM, or box-plots showing median (solid line), mean (dotted line) and 10^th^, 25^th^ and 75^th^ and 90^th^ percentiles. * represents main effect of GENOTYPE *p*<0.05; ## represents main effect of within-subject factor (TRIAL, PREPULSE, SESSION or SPEED).

As expected, increasing the prepulse volume led to a linear increase in the inhibition of the startle in both groups (Figure 1B; ANOVA, main effect of PREPULSE, F_1.49,80.35_=73.26, *p=*0.001). However, there was a 20-40% PPI reduction in the *Ehmt1*^D6Cre/+^ mice relative to *Ehmt1*^fl/+^ cage-mates (Figure 1B; ANOVA main effect of GENOTYPE, F_1,54_=5.54, *p*=0.022), suggesting that, in addition to an enhanced startle response, the mutant mice were also impaired in the normal PPI response

### Normal motor function in the *Ehmt1*^D6Cre/+^ mouse

Altered sensory-motor gating in *Ehmt1*^D6Cre/+^ mice was not due to any gross deficits in motoric competence. Training on the rotarod test indicated normal learning with repeated sessions, with latency to fall reducing across training sessions (Figure 1C; ANOVA, main effect of SESSION, F_5,130_=21.33, *p*<0.001), but there was no difference in GENOTYPE (F_1,26_=0.12, *p*=0.73) and no interaction between training SESSION and GENOTYPE (F_5,130_=0.67, *p*=0.65). Moreover, in an accelerating rotarod probe test of motor coordination, latency to fall decreased as the speed increased (Figure 1D; ANOVA, main effect of SPEED, F_4,104_=38.38, *p*<0.001), but there was no difference between GENOTYPE (F_1, 26_=0.67, *p*=0.42) or interaction between SPEED and GENOTYPE (F_4,104_=0.65, *p* =0.63).

### Reduced anxiety in the *Ehmt1*^D6Cre/+^ mouse

A number of measures in the elevated plus maze and open field tests indicated that *Ehmt1*^D6Cre/+^ mice have a reduced anxiety phenotypes. On the EPM (Figure 2A and B) *Ehmt1*^D6Cre/+^ mice spent on average twice as long as *Ehmt1*^Fl/+^ cage-mates on the open-arm (*t*_*26*_ =-2.08, *p* = 0.04) and made on average 40% more entries into the open arm of EPM more than controls, although this did not reach statistical significance (*t*_*26*_ = -1.52, *p =* 0.14). On the OF test (Figure 2C and D), *Ehmt1*^D6Cre/+^ mice made on average 25% more entries in the inner zone of the OF (*t*_*26*_= -3.21, p = 0.004) and spent 50% more time in the inner zone than *Ehmt1*^Fl/+^ mice, although this did not reach statistical significance (*t*_*26*_=-1.86, *p*=0.07).

**Figure 2.**
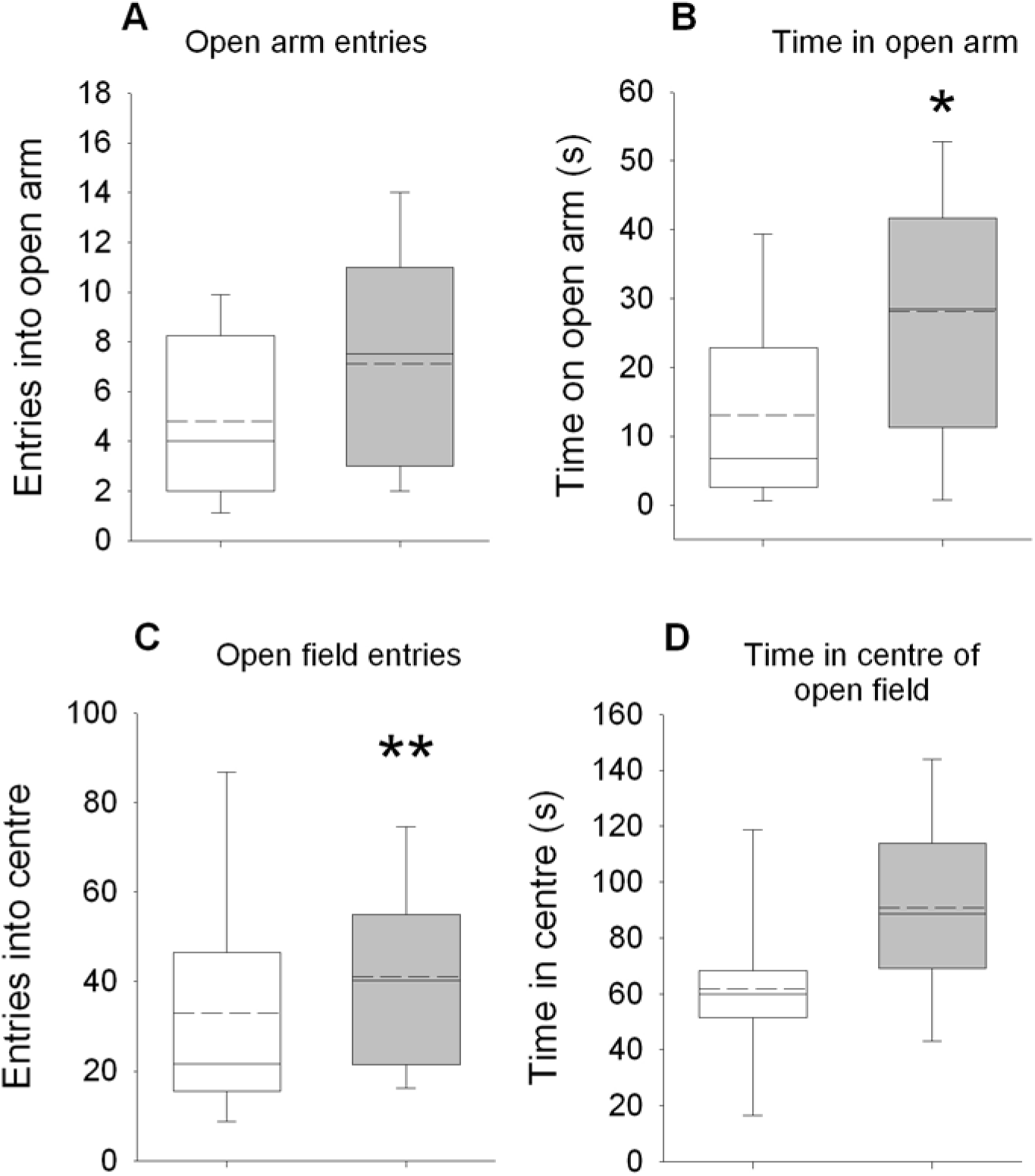
Behaviour in the elevated plus maze and open field. On the EPM *Ehmt1*^D6Cre/+^ mice (n=17) showed a pattern of behaviour consistent with reduced anxiety relative to *Ehmt1*^Fl/+^ mice (n=9), including increased open arm entries (**A**) and increased time on time on the open arm (**B**), although only the latter was statistically different from controls. Convergent evidence of a reduced anxiety phenotype was seen in the OF test. Here *Ehmt1*^D6Cre/+^ mice made significantly more entries into the centre of the OF (**C**) and spent more time in the inner zone (**D**), although this did not reach significance. Box-plots showing median (solid line), mean (dashed line) and 10^th^, 25^th^ and 75^th^ and 90^th^ percentiles; * represents main effect of GENOTYPE *p*<0.05; ** *p*<0.01.

### Impaired novel object recognition in *Ehmt1*^D6Cre/+^ mouse

The novel object recognition (NOR) task takes advantage of the preference of rodents to attend to new objects in their environment as a means for testing declarative memory (Antunes and Biala, 2012). Here, two inter-trial intervals were used: half of the cohort was examined following a 15-minute delay between the initial object exposure and the test trial, and half the cohort was examined after a 90-minute delay. In the test phase, as expected the *Ehmt1*^fl/+^ control mice explored the novel object significantly more than chance (50%) in both the 15-minute and 90-minute retention trials (Figure 3; Kolmogorov Smirnov test (KS), 15min, 60%, *p*=0. 02; 90min retention trial, 67%, *p*=0.002). However, exploration of the novel object by *Ehmt1*^D6Cre/+^ mice was not significantly above chance, at either 15min (52%, *p=0.*97), or 90 min retention trial (50%, *p*=0.44).

**Figure 3.**
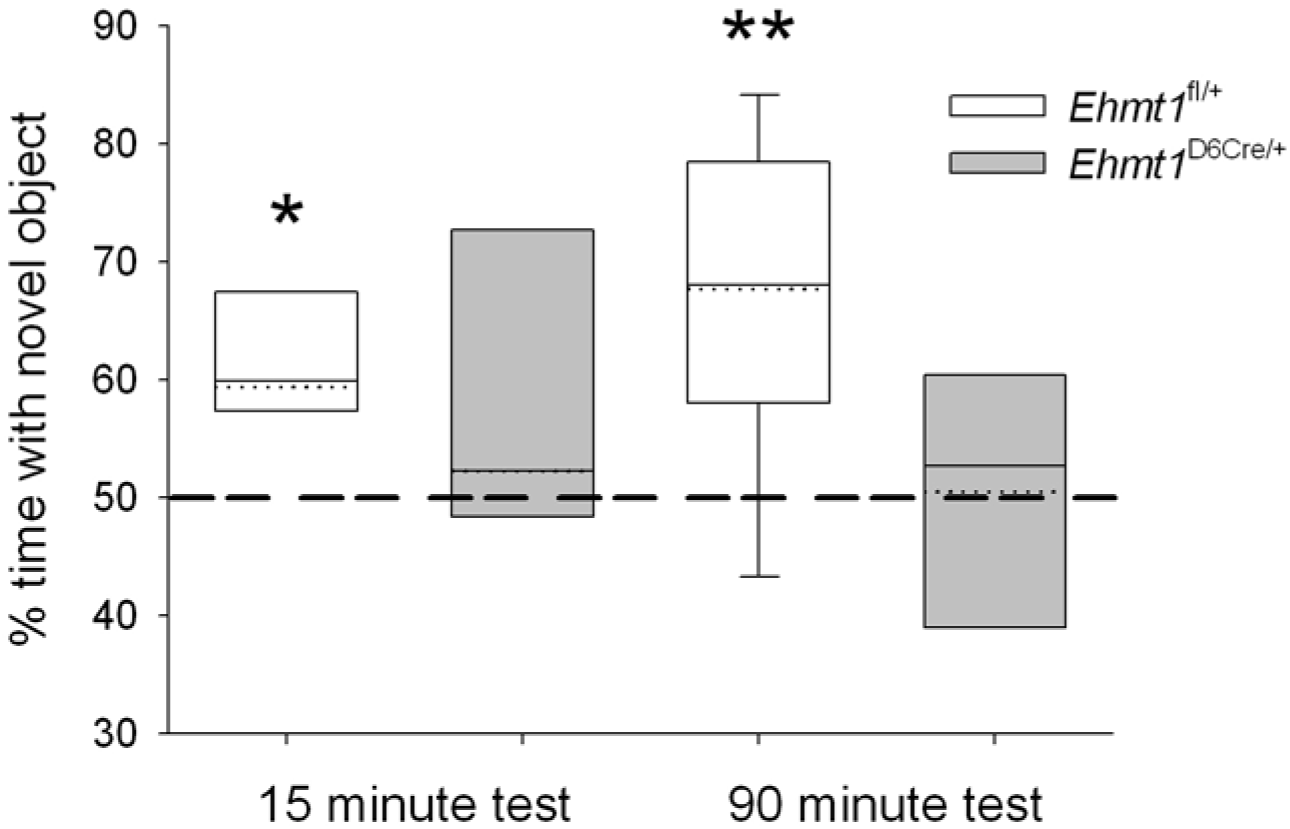
Novel object recognition. In both the 15-minute and 90-minute retention tests of the NOR task *Ehmt1*^fl/+^ control mice (n=16) explored the novel object more than chance. However, exploration of the novel object by *Ehmt1*^D6Cre/+^ mice (n=13) was not significantly above chance, for either the 15- or 90-minute retention trials indicating a deficit in declarative memory. Data are box-plots showing median (solid line), mean (dotted line) and 10^th^, 25^th^ and 75^th^ and 90^th^ percentiles (dots represent 5^th^ and 95^th^ percentile). * represents main effect of GENOTYPE *p*<0.05; ** *p*<0.01.

This difference at test was not due to differences during the habituation phase, as indicated by overall exploration time of the object that was replaced by a novel object at test (*Ehmt1*^fl/+^ mean=114s, ±SEM=15; *Ehmt1*^D6Cre/+^ mean=113s ±SEM=12; ANOVA, no main effect of GENOTYPE, F_1,28_=0.031, *p*=0.86).

### Electrophysiological measurements of paired pulse AEPs

In order to gain an insight into the neural changes underpinning the deficits in sensory-motor gating seen in *Ehmt1*^D6Cre/+^ mice we used electrophysiological methods. A subset of the behavioural cohort was then subject to EEG recording to measure auditory evoked potentials (AEPs). AEPs are voltage fluctuations time-locked to an auditory event used for brain dysfunction clinical diagnosis (Luck et al., 2011). We used a paired pulse paradigm in which two pulses were delivered back-to-back with a 500ms interval between the first stimulus (S1) and the second stimulus (S2) (Figure 4A). The grand average waveforms show a stereotypic maximal positive deflection (P1) and maximal negative deflection (N1) (Figure 4B1).

**Figure 4.**
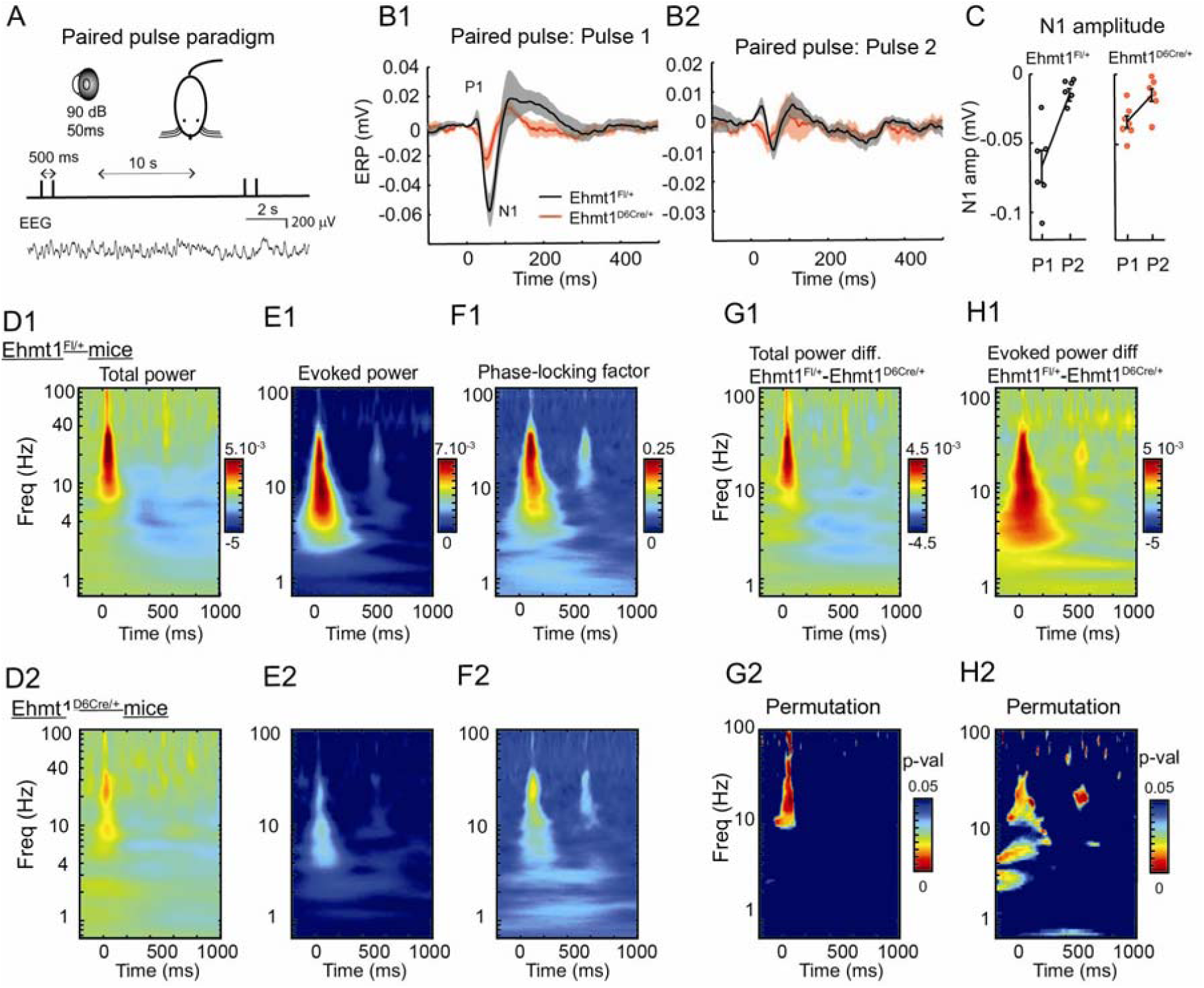
AEP paired pulse measurements. **A**) Schematic showing the paired pulse stimulus paradigm. **B1**) Grand average ERP waveform after pulse 1 (S1) in *Ehmt1*^D6Cre/+^(red) *and Ehmt1*^fl/+^ (black). The P1 and N1 components are indicated on the waveform. **B2**) The waveforms for pulse 2 (S2); **C)** N1 amplitude for *Ehmt1*^D6Cre/+^ (n=7) and *Ehmt1*^fl/+^ (n=7) mice. **D1 & D2)** Normalized total power (in time-frequency plot) of both S1 (time 0) and S2 (time 500ms) for *Ehmt1*^fl/+^ (D1) and *Ehmt1*^D6Cre/+^ (D2). Note the early increase in the beta/gamma range (10-100Hz) and the late decrease in the delta range (∼4Hz) and the reduction of both components in the *Ehmt1*^D6Cre/+^. **E1 & E2)** Evoked power time-frequency plots for the same data set; **F1 & F2)** Phase locking factor for the same data set; **G1)** Difference of total power between *Ehmt1*^fl/+^ and *Ehmt1*^D6Cre/+^. **G2)** Statistical significance heat map based on permutation tests of total power are indicated by the colour scale (*p*-value < 0.05); **H1)** Difference of evoked power between *Ehmt1*^fl/+^ and *Ehmt1*^D6Cre/+^ mice; **H2)** Statistical significance between evoked power of *Ehmt1*^fl/+^ and *Ehmt1*^D6Cre/+^ mice (spurious spots above 50Hz are due to occasional 50Hz noise contamination in some of the recordings). Data shown is mean ±SEM.

*Ehmt1*^D6Cre/+^ mice had a nearly two-fold lower N1 amplitude response after the S1 (Figure 4B1) but not after S2 (Figure 4B2). Additionally, *Ehmt1*^D6Cre/+^ mice had a significant reduction in the S2/S1 ratio for the N1 component (Wilcoxon, rank sum=64, *p*=0.018), what is considered an electrophysiological measurement of sensory gating (Figure 4C). During peak activation of the paired pulse experiment we observed an increase in total power across high frequency bands from 10-100Hz in both groups of mice (Figure 4D1 and D2). The difference time-frequency plot and the permutation test between *Ehmt1*^D6Cre/+^ and *Ehmt1*^fl/+^ control mice however, revealed that the distributed peak that occurred approximately 40ms after the S1 pulse, a time corresponding with the N1, was significantly greater in the beta (13-30Hz) and gamma (30-70Hz) frequency bands in the *Ehmt1*^fl/+^ control mice (Figure 4G1 and G2). The late (>200ms after the S1 pulse) decrease in the delta frequency band (∼4Hz) prominent in *Ehmt1*^fl/+^ (Figure 4D1-G1) was not different from *Ehmt1*^D6Cre/+^ (Figure 4G2).

Measurements of evoked power, EEG power which is phase-locked with the event onset across trials, demonstrated increases in the delta (∼4hz), beta (13-30Hz) and low gamma (here ∼40Hz) band responses in both groups of mice approximately ∼30-50 ms after the S1 (Figure 4E1 and E2). Again, permutation tests revealed a reduction in evoked power in *Ehmt1*^D6Cre/+^ mice after both the S1 and ∼40ms after the S2 pulse (Figure 4H1 and H2). In complement, we measured the phase locking factor (PLF), which provide a measurement of trial-to-trial reliability across the frequency domain (Goffin et al., 2014). To extract the PLF, magnitude information is transformed to 1 and averaged so that the phase angle with respect to event onset is extracted (Roach and Mathalon, 2008). Values between zero and one are obtained, in which one reflects perfect synchrony of phase angles across trials. *Ehmt1*^D6Cre/+^ mice did not show PLF values above .17 at any point, while control mice show nearly two-fold PLF synchrony (+.25) at ∼40ms post-S1 pulse, between ∼20-40Hz. Overall *Ehmt1*^D6Cre/+^mice demonstrated reduced PLF (Figure 4F1 and F2).

### Electrophysiological measurements of Mismatched Negativity AEPs

The Mismatch Negativity (MMN) response is elicited when a qualitative feature of the stimulus does not match the pattern in a previous series (Light and Braff, 2005) (Figure 5A). One core feature of MMN is the importance of NMDA receptor function. For instance, non-competitive NMDA-R antagonists, like ketamine and the selective antagonist of the NR2B NMDA subunit, CP-101,606 reduce MMN amplitude (Sivarao et al., 2014), the level of which predicts the magnitude of psychotic experiences in response to these drugs (Umbricht et al., 2002). Therefore, in order to probe this function further here, mice were given either saline or 10mg/kg of ketamine prior to the MMN test.

**Figure 5.**
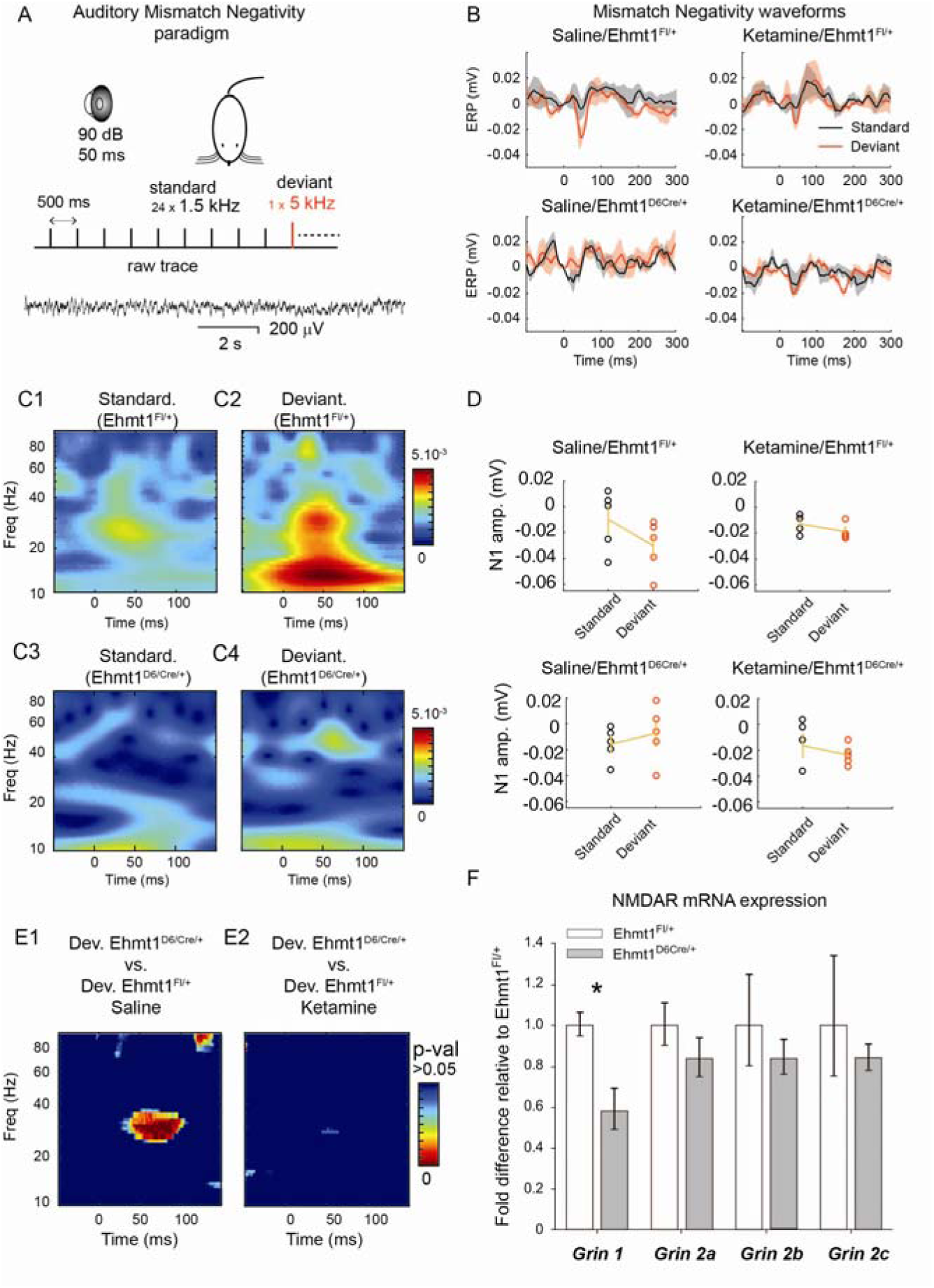
AEP mismatch negativity phenotypes & NMDA mRNA expression. **A)** Mismatch negativity stimulus **B)** Grand average ERP waveforms (shaded areas) after the saline or ketamine conditions in the in *Ehmt1*^fl/+^ (n=5) and *Ehmt1*^D6Cre/+^ (n=5), for the deviant (red) and standard (black) tone for each condition. **C1 & C2)** Time-frequency plot of evoked power for the standard and deviant pulse in *Ehmt1*^fl/+^ mice after saline injection; **C3 & C4)** Time-frequency plot of evoked power for the standard and deviant pulse in *Ehmt1*^D6Cre/+^mice following saline injection. **D)** N1 amplitude peak amplitude corresponding to each condition. **E1)** Permutation test showing heat map of *p*-values for the difference of distribution between deviant tone maps of saline treated *Ehmt1*^fl/+^ (C2) and *Ehmt1*^D6Cre/+^ (C4). **E2**) Permutation test showing heat map of *p*-values for the difference of distribution between deviant tone maps of ketamine treated *Ehmt1*^fl/+^ (C2) and *Ehmt1*^D6Cre/+^ (C4). **F)** NMDA-R subunit mRNA expression in hippocampal samples (*Ehmt1*^fl/+^ n=4, *Ehmt1*^D6Cre/+^ n=8). Data shown is mean ±SEM. * represents main effect of GENOTYPE *p*<0.05.

In saline treated animals there was a difference between *Ehmt1*^D6Cre/+^ and *Ehmt1*^*fl/+*^ mice in response to the deviant pulse (Figures 5C-D) as indicated by analysis of the N1 amplitude that showed an interaction between GENOTYPE and PULSE (repeated measures ANOVA, F_1,8_=12.70, *p*=0.007). In the standard pulse condition *Ehmt1*^*fl/+*^ mice, amplitude corresponded to an increase in ∼10-40Hz evoked potential approximately 30-50ms post-stimulus (Figure. 5C1-C2). In the deviant pulse condition, *Ehmt1*^*fl/+*^ mice not only show an increase in ∼10-40Hz evoked potential, but also one with greater peak latency, at 40-70ms post-stimulus (Figure 5C2). In contrast, *Ehmt1*^D6Cre/+^ mice treated with saline showed no pattern of change in the amplitude response for either standard or deviant pulses.

However, following 10mg/kg of ketamine administration, the difference in the N1 amplitude between the standard and deviant pulse was absent for both genotypes (Figure 5B and D), as indicated by a lack of interaction between GENOTYPE and PULSE (repeated measures ANOVA, F_1,7_=0.011, *p*=0.921). Further analysis underlined this finding. Permutation tests after saline administration for the deviant-only condition demonstrates a difference between *Ehmt1*^*fl/+*^ controls and *Ehmt1*^D6Cre/+^ wavelet transforms, at 30Hz between 50-80ms post-stimulus (Figure 5E1), driven by the higher amplitude response in the *Ehmt1*^*fl/+*^ control mice. In contrast, after ketamine administration the difference in deviant response between *Ehmt1*^*fl/+*^ controls and *Ehmt1*^D6Cre/+^ mice was abolished (Figure 5E2).

### Altered NMDA-receptor expression in *Ehmt1*^D6cre/+^ mutant mice

In light of similarity in basal *Ehmt1*^D6Cre/+^ electrophysiological phenotype and the electrophysiological phenotype in *Ehmt1*^*fl/+*^ control mice following ketamine administration, we assessed the NMDA system in *Ehmt1*^D6Cre/+^ mice. RT qPCR was used to examine the mRNA levels of NMDA-R subunits *Grin1, Grin2a, Grin2b* and *Grin2c* in the adult hippocampus. Expression of *Grin1*, the gene encoding NMDA NR1 subunit, was significantly reduced by 40% on average in *Ehmt1*^D6cre/+^ mice (Figure 5F; t=-3.07, *p*=0.014). No difference was seen in expression of other NMDA-R subunits examined (Figure 5F *Grin2a*, t=-1.02, *p*=0.331; *Grin2b*, t=-0.814, *p*=0.435; or *Grin2c*, t=-0.71, *p*=0.498). Furthermore, there were no genotype differences in AMPA-R subunit gene expression examined (See Supplementary Figure 4).

## DISCUSSION

Recent studies in human genomics strongly implicate a number of genes found to regulate chromatin dynamics in the etiology of neurodevelopmental disorders from developmental delay (Singh et al., 2016), to autism spectrum disorders (RK et al., 2017, De Rubeis et al., 2014), and schizophrenia (Network and Pathway Analysis Subgroup of Psychiatric Genomics, 2015). *EHMT1* is one such gene, associated with both neurodevelopmental and neuropsychiatric disorders. Here we examined behavioural and electrophysiological correlates of information processing in a mouse model of *Ehmt1* haploinsufficiency, specifically in the forebrain. We found that *Ehmt1*^D6Cre/+^ mice have sensorimotor and auditory gating deficits, reduced anxiety, and learning and memory deficits, in the absence of generic deficits in motor function. *Ehmt1*^D6Cre/+^ mice also showed a number of abnormalities in electrophysiological measurements including a reduced magnitude of AEPs after paired-pulse inhibition and MMN tasks, reduced evoked and total power in high frequency bands, and reduce PLF. Overall these data indicate that *Ehmt1*^D6Cre/+^ mice show deficits in sensory motor gating and information process, possibly related to abnormal NMDA-R functioning.

*Ehmt1*^D6Cre/+^ mice displayed decreased anxiety in a number of measures across a two separate tests, namely the EPM and OF. Although these findings in our model were not confounded by impaired motor competence as indicated by normal behaviour and learning on the rotarod test, not all measures on both the EPM and OF reached statistical significance. It may be that a more robust assessment of anxiety in this model would be achieved using a unified score from a number of separate measures (Harrison et al., 2020). Generally however, these findings of decreased anxiety are consistent with previous studies of *Ehmt1* haploinsufficient models (Iacono et al., 2018, Balemans et al., 2010), although other models, specifically a CamKII-drive full *Ehmt1* deletion shows decreased anxiety (Schaefer et al., 2009). This indicates that the relationship between *Ehmt1* function and anxiety behaviour probably depends on extent, timing and location of the genetic lesion.

*EHMT1* is a risk gene associated with a number of neurodevelopmental and psychiatric disorders (Talkowski et al., 2012), thus it was important for this work to focus not only on translational phenotypes but those that overlap traditional diagnostic boundaries. Several prominent features shared by these disorders are associated with deficits in attention and gating, or filtering out intrusive stimuli (Javitt, 2009, Orekhova et al., 2008, Perry et al., 2007, Belmonte et al., 2004). Behaviorally, the *Ehmt1*^D6Cre/+^ mice showed evidence of this in an acoustic startle test. Consistent with previous findings in other *Ehmt1* haploinsufficiency models (Iacono et al., 2018, Balemans et al., 2013), sensory motor gating deficits were manifest by both a greatly enhanced acoustic startle response, and a decreased pre-pulse inhibition of startle.

Information-processing deficits are another common phenotype associated with *EHMT1* risk populations. Accordingly, *Ehmt1*^D6Cre/+^ mice were examined on their performance in the NOR task, a paradigm where an animal’s ability to remember a previously encountered object is indicated by an increased willingness to explore the novel object over the familiar object. The *Ehmt1*^D6Cre/+^ mice showed no evidence of novel object recognition, even with only a 15-minute interval after habituation. Importantly, this was not due to reduced interest in exploration or exposure to the familiar object, as time spent in exploration was equivalent across *Ehmt1*^fl/+^ control and *Ehmt1*^D6Cre/+^ mutant mice. Similarly to the acoustic startle and prepulse inhibition phenotype, deficits in a NOR paradigm have been seen in other *Ehmt1*^*+/-*^ mice (Iacono et al., 2018, Balemans et al., 2013). This convergence across models and laboratories suggests that both sensory motor-gating and information processing deficits are key behavioural features of *Ehmt1* haploinsufficiency.

Given the robustness of behavioural deficits in sensory motor-gating and information processing seen in models of *Ehmt1* haploinsufficiency, we explored this further by examining electrophysiological measures of these domains. *Ehmt1*^D6Cre/+^ had reduced N1 amplitude in response to stimulus 1 (S1). Furthermore, *Ehmt1*^D6Cre/+^ mice had a significant reduction in the S2/S1 ratio for the N1 component, an electrophysiological correlate of sensory gating.

Evidence of information-processing deficits in the *Ehmt1*^D6Cre/+^ mice was provided by the reduced response in a mismatch negativity (MMN) paradigm. The decreased amplitude responses in these AEP measurements meanwhile corresponded to significant reductions in total and evoked power in high frequency bands. Such reductions may be particularly insightful, as numerous studies report disruptions in gamma-band activity across neurodevelopment (Orekhova et al., 2008, Kwon et al., 1999), neuropsychiatric (Tatard-Leitman et al., 2015) and neurodegenerative diseases (Iaccarino et al., 2016). Whether reduced gamma activity is actually directly associated with disease pathologies across these patient populations remains largely unknown. Recently however, reductions in gamma-band activity were found to precede the onset of plaque formations and cognitive decline in a mouse model of Alzheimer’s disease.

Meanwhile the stimulation of fast-spiking interneurons in the gamma frequency range (∼40Hz), a way to boost gamma-band activity (Cardin et al., 2009), led to the reduction in the plaque forming amyloid isoforms and attenuated plaque load in ageing mice (Iaccarino et al., 2016). In a more functional assessment, the induction of long-term potentiation using high frequency stimulation (∼100Hz) was found to restore spine density and long-term memory in early stages of the disease in mice (Roy et al., 2016). While these findings are specific to Alzheimer’s disease, they also confirm an important link between gamma synchrony and cognitive function exists (Fries, 2015). Furthermore, we may find disruptions in high frequency oscillation patterns tightly correspond with the degree of cognitive impairment and range of pathologies across *EHMT1* risk populations.

The decreases in high frequency activity coupled with reduced *Grin1* mRNA are suggestive of overall disruptions in local connectivity and may hint to more global imbalances in excitation/inhibition (E/I). Such findings do corroborate previous reports showing abnormalities in neural development and connectivity in *Ehmt1*^+/-^ mouse models (Benevento et al., 2017, Balemans et al., 2014, Balemans et al., 2013), and that *Ehmt1-*mediated H3K9me2 levels dynamically regulate synaptic scaling, thus playing a direct role in the fine balance between excitation and inhibition at the level of individual neurons (Benevento et al., 2016). And indeed many of the *Ehmt1*^D6Cre/+^ mice phenotypes are markedly similar to *Grin1* mouse mutants (Furuse et al., 2010). The *Grin1*^*neo-/-*^ mice and the *Grin1*^*Rgsc174*^ heterozygous mice both show increased stereotypy (Mohn et al., 1999) and deficits in sensorimotor gating (Duncan et al., 2004). Mice with even subtle reductions in the NR1 receptor are found to have decreases in MMN (Featherstone et al., 2015), gamma-band disruptions and reduction in E/I balance (Gandal et al., 2012). Nevertheless, it is important to note that the studies presented here represent a change in mRNA and not protein levels and, moreover, only provide a correlative link between the electrophysiological deficits seen in *Ehmt1*^D6Cre/+^ mice and *Grin1* reduction. Interestingly however, a recent study has demonstrated *increased Grin1* expression and NMDA-R hyperactivity in iPS cells derived from Kleefstra syndrome patients (Frega et al., 2019). Despite the discrepancies, together these data suggest that future investigation of *Ehmt1* haploinsufficiency may benefit from further examination of the relationship with the NMDA system.

In summary, *Ehmt1*^D6Cre/+^ forebrain-specific haploinsufficiency produced deficits in sensory-gating and information processing. Behavioural evidence from explicit tests of sensorimotor gating and findings from a learning and memory task, suggest that *Ehmt1*^D6Cre/+^ mice do not attend to or process information in order to inform appropriate behavioural responses. Neural correlates of these abnormalities were further demonstrated using electrophysiological studies, which indicated deficits related to disruptions in local connectivity and NMDA function. Taken together these data suggest that *Ehmt1* haploinsufficiency leads to abnormal circuit formation and behavioural abnormalities that likely underpin deficits seen across the broad spectrum of neurodevelopmental and neuropsychiatric disorders with which *EHMT1* is associated.

## Data availability

Much of the data presented in this paper are provided as supplementary files. Any materials that are not, will be made available by the authors on request where possible.

## Author contribution statement

*Conceptualization* BAD, FD, ARI; *Data curation* BAD, FD, CO’R, MA; *Formal Analysis* BAD, FD, ARI; *Funding Acquisition* BAD, ARI; *Investigation Methodology* BAD, FD; *Project Administration* BAD, FD, ARI; *Resources* VC, ARI; *Software* BAD, FD; *Supervision* AJH, VC, ARI; *Validation* BAD, FD; *Visualization* FD, MA, ARI; *Writing – original draft* BAD, FD; *Writing – review & editing* BAD, FD, ARI.

