## Supplementary for "Impairments in sensory-motor gating and information processing in a mouse model of *Ehmt1* haploinsufficiency"

### – Supplementary Materials

Brittany A. Davis[1], François David[1], Ciara O'Regan[2], Manal Adam[2], Adrian J. Harwood[1], Vincenzo Crunelli[1] & Anthony R. Isles[2*]

[1]Neuroscience and Mental Health Research Institute and School of Biosciences, Cardiff University, Cardiff, UK

[2]MRC Centre for Neuropsychiatric Genetics and Genomics, School of Medicine, Cardiff University Cardiff, UK

**Running title: *Ehmt1* and information processing**

**RESULTS**

#### Supplementary Figure 1

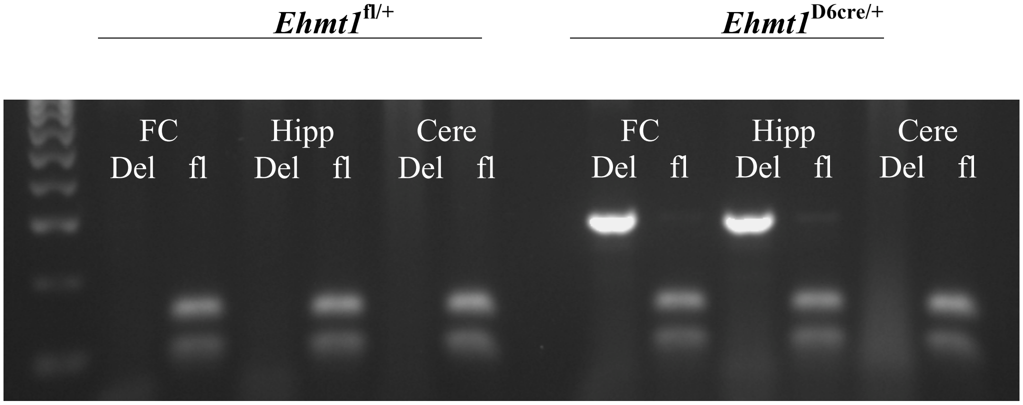


**Supplementary Figure 1. Confirmation of selective deletion of Ehmt1 under the D6-cre transgene.** PCR analysis demonstrated that the deleted allele (“Del” lanes) was present in frontal cortex (FC) and hippocampus (Hipp), but not cerebellum (Cere) of *Ehmt1*$^{D6cre/+}$ mice only. PCR for fl and wild-type *Ehmt1* (“fl” lanes) indicated the presence of DNA in all samples.

#### Supplementary Figure 2

**A** **B**

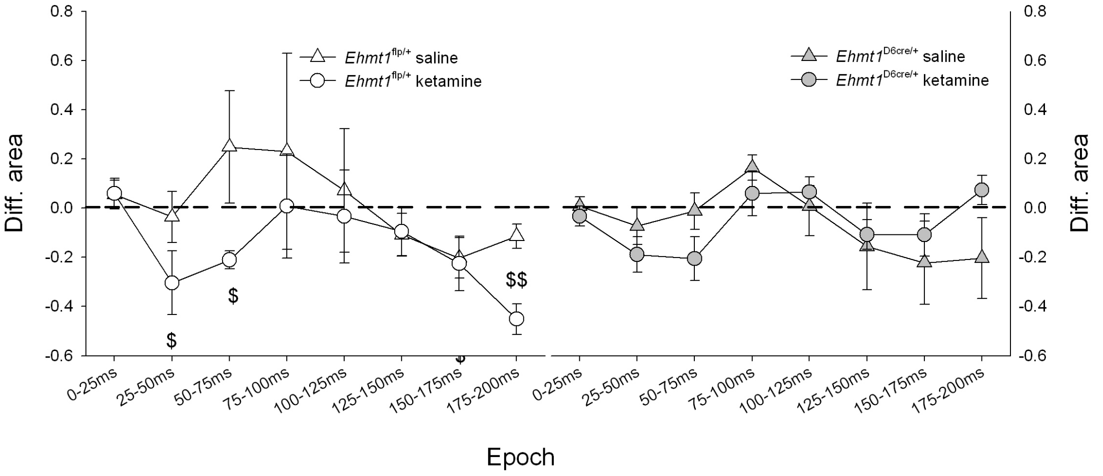


**Supplementary Figure 2 Epoch analysis in *Ehmt1*$^{D6cre/+}$ and control mice** A) MMN area wavelet differences (Standard condition subtracted from Deviant condition) across 8 time epochs from 0-25ms to 175-200ms, for the *Ehmt1*$^{fl/+}$ after administration of saline (triangles) and ketamine (circles). Above zero values indicate larger wavelet epochs in the deviant condition, while below zero values indicate larger wavelet epochs in the standard condition. Only epochs with greater than zero values were found in the saline condition, after the classic N1 time epoch (25-50ms) at the 50-75ms and 75-100ms epochs, suggesting a delayed N1 or the MMN response. These values did not reach significance, due to level of variability. After ketamine administration we find the classic N1 time epoch 25-50ms as well as 50-75ms were significantly different from zeros. However, these values were below zero, thus the standard condition wavelets were larger than deviant condition suggesting the predicted loss of MMN detection after ketamine administration occurred in controls. B) MMN area wavelet differences across 8 time epochs for the *Ehmt1*$^{D6cre/+}$ mutant mice after administration of saline (triangles) and ketamine (circles). There was a slightly above 0 value in mutant mice at the 75-100 epoch, however overall at no epoch was there a significant difference from 0 and little to no change in the overall pattern of response was seen between the ketamine and saline condition in these mice. Data are means ± SEM; $ represents significant difference in ketamine condition ($p<0.05$; $$, $p<0.01$)

#### METHODS

##### PCR for deletion specifcity

To establish *Dach1-Cre* specificity and verify the accuracy of the PCR methods for genotyping, sections from the prefrontal cortex (PFC) and the hippocampus were taken for positive verification of the deleted allele in the mutants (985-bp band in the gel). While a section of the cerebellum, a region where *Dach1* is not expressed was taken to check for the non-deleted floxed allele (792-bp). A positive control for the wild type allele was included (95-bp). In addition sections from control PFC, hippocampus and cerebellum animals, which are expected to have one copy of the floxed-allele, were also examined. PCR primers for *Ehmt1*$^{fl/-}$Forward (5-GCCTGGTGAATTTTAGTGGGC-3); Reverse 1 (5-GTTTGGGGCAAGTGTGGAG-3) Reverse 2 (5-TTGTACAAGAAAGCTGGGTCT-3). The PCR profile used was 94°, 5min, 94°, 40s, 60°, 45s, 72°, 1min for 35 cycles and 72°, 10min.

##### Electrode implantation

Mice were anesthetized with 2% isoflurane and underwent stereotaxic surgery for the implantation of five enameled stainless steel electrodes (diameter 70mm, California Fine Wire Company): two bilateral frontal electrodes, one monopolar and one bipolar (2.7-mm anterior, 1.5-mm lateral, 1.2-deep relative to bregma); two bilateral hippocampal electrodes, one monopolar and one bipolar (2.7 mm posterior, 3-mm lateral, 2.2-mm deep relative to bregma); and one bipolar electrode in the auditory cortex (2.7-mm posterior, 4-mm lateral, 1.1-deep relative to bregma) (Supplementary Figure 1). The two monopolar electrodes were implanted on the right hemisphere and differential signals between these two electrodes were recorded. Bipolar electrodes were implanted in the left hemisphere. Two epidural screw electrodes were placed above the cerebellum to ground the animal. All electrodes were connected to a Precidip (Switzerland) connector and secured with the use of superglue and dental cement (acrylic and Metabond cement (Sun Medical Co., Japan)). Animals were allowed 7-days recovery before recordings took place. All data reported here are from the differentially recorded signal between the right frontal and hippocampal monopolar electrodes. Previous studies demonstrate that this electrode configuration produces AEP components most characteristically similar to human EEG central cortical scalp recordings (Siegel *et al.*, 2003; Ehrlichman *et al.*, 2008).

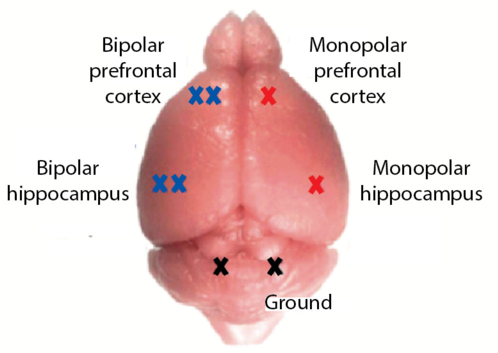


**Supplementary Figure 3 Stereotaxic placement of electrodes**

Five wire electrodes: three subdural electrodes were placed in the left auditory cortex auditory, and right and left hippocampus; and two supradural, one in the right and left neocortex. Two screw electrodes were placed in the cerebellum for ground.

#### EEG recording & analysis

The animal connector was connected to a SuperTech Bio-AMP(Pecs) pre- (bandwidth 0.1–500 Hz) and main-amplifiers (band-width DC to 500 Hz), the signal acquired at 1kHz using a Cambridge Electronic Design, CED(UK), Micro3 D.130 (1401) digitizer and CED Spike2 v7.3 software.

Auditory stimuli (50ms sinusoid at 1.5 and 3kHz) were generated using the same software and were delivered through a Power1401 interface (CED). Auditory stimuli were delivered with speakers positioned directly in front of each recording cage. The sound pressure was calibrated at 90dB by using a sound meter based on the approximate height of the animal's head within the plexiglass cage. Mice were given 30mins to acclimate for the first 2 recording trials and 15-mins thereafter. Mice were recorded in the dark, and during the dark cycle in order to record during their normal active state.

Data analysis was performed using MatLab (MathWorks, Natick, MA) software. Due to the loss of EEG signals in later recording sessions, some mice were excluded from the analysis.

ERPs were obtained by averaging epochs centred at Time 0 and 500msec to 0mV, respectively. For each epoch, power was calculated using either an fft or wavelet transform. For the wavelet transform, EEG signals were transformed using the complex Morlet's wavelets $w(t, f_0)$(Kronland-Martinet *et al.*, 1987). The script used was the wt.m found at https://www.physics.lancs.ac.uk/research/nbmphysics/diats/tfr/. The wavelets have a Gaussian shape in time (Standard deviation $\sigma_t$) and frequency (Standard deviation $\sigma_t$) around its central frequency $f0$: $w(t, f_0) = A * exp(-t^2/2\ \sigma_t^2) * exp(2i\pi f_0 t)$ with $\sigma_f = 1/\pi\sigma_t$ (Tallon-Baudry et al., 1996). All the wavelet used had a constant ratio $f_0/\sigma_f = 1$, with $f_0$ ranging from 0.5 to 100Hz in logarithmically distributed frequency steps.

The wavelet transform produces a time-frequency representation of power (TF energy). Measurements can be applied to either the averaged evoke potential or to the individual trials, thus generating either evoked power or total power, respectively. For total power, the noise remains in the signal, which may mask any activity that does not have a high signal-to-noise ratio. Thus for total power a baseline correction is applied. The baseline pre-stimulus period from –600 to –200 was subtracted from the entire peri-stimulus period and applied to the frequency bands independently.

In order to extract the phase-locking factor of the oscillatory burst the normalized complex time-varying power of each trial was averaged (Tallon-Baudry *et al.*, 1996). By averaging, complex values are produced, which includes phase information at each time-frequency region around $t$ and $f_0$. The phase-locking factor is the calculated by unit normalizing or transforming the magnitude information of the complex value. The value remaining represents phase information and is an integer from 0 to 1, which ranges from non-phase-locked or 0, to strictly phase locked or 1. Such methods are found to be robust against artefacts.

#### Permutation testing

Statistical tests on the frequency data was performed using the permutation method (Westfall and Young, 1993), with the maximal number of iterations allowed by the compared samples or 1,000 iterations in case it surpasses this number. The number of iterations was dependent on the total number of possible permutations (which varied with the number of animals included in analysis). In the paired-pulse analysis, t-tests were calculated in between controls and mutants around the first pulse. In the MMN analysis, t-tests were calculated in the mutant between $24^{th}$ pulse and the deviant pulse; and in the control between $24^{th}$ pulse and the deviant pulse. Separate permutation plots were generated for the saline and ketamine condition. In addition, t-tests were calculated between mutant and controls for the ketamine condition and for the saline condition. Statistical permutation plots were generated in all of the above conditions for total power and evoked power.
